# A conserved tyrosine in the S1/S2 loop shapes SARS-CoV-2 spike processing and host protease usage

**DOI:** 10.64898/2026.11.12.766136

**Authors:** William M. Meyers, Angelica L. Morgan, Michelle N. Vu, Kumari G. Lokugamage, Yani P. Ahearn, Leah K. Estes, Yiyang Zhou, R. Elias Alvarado, Mehul S. Suthar, David H. Walker, William K. Russell, Bryan A. Johnson, Vineet D. Menachery

**Affiliations:** Experimental Pathology Graduate Program, University of Texas Medical Branch, Galveston, Texas, USA; Department of Microbiology and Immunology, University of Texas Medical Branch, Galveston, Texas, USA; Department of Pediatrics, Emory University School of Medicine, Atlanta, Georgia, USA; Emory Vaccine Center, Emory University, Atlanta, Georgia, USA; Department of Microbiology and Immunology, Emory University School of Medicine, Atlanta, Georgia, USA; Department of Pathology, University of Texas Medical Branch, Galveston, Texas, USA; Center for Biodefense and Emerging Infectious Disease, University of Texas Medical Branch, Galveston, TX, United States; Department of Biochemistry, University of Texas Medical Branch, Galveston, Texas, USA

**Author notes:** Corresponding Author: Vineet D. Menachery, **Email:**.

## Abstract

Both spike processing efficiency and host protease usage play critical roles in SARS-CoV-2 infection, pathogenesis, and transmission. The SARS-CoV-2 spike contains two motifs, a furin cleavage site (FCS) and an extended S1/S2 loop, which improve both cleavage efficiency and host protease usage. Disruption of these motifs significantly attenuates SARS-CoV-2 infection and pathogenesis. Importantly, modifications and changes within the S1/S2 loop can also shape infection and disease outcomes following infection. In this study, we examined mass spectrometry data generated from purified virions from wild-type (WT) or mutant SARS-CoV-2 infection. We identified phosphorylation occurring at highly conserved residue Y674 and used reverse genetics to generate a mutant that ablates phosphorylation capacity in the S1/ S2 loop (Y674F). Our results show that the Y674F mutant replicated equivalently to wild-type SARS-CoV-2 in Vero cells but had attenuated replication in a human respiratory cell line. Golden Syrian hamsters infected with the Y674F mutant had equivalent weight loss, lung viral titer, and similar immune pathology to WT SARS-CoV-2. However, the mutant had reduced viral load in the nasal wash and reduced antigen staining in the large airways in the lung. Notably, loss of Y674 ablated spike processing to the S1/S2 cleavage product on virions. In addition, Y674F mutant had a diminished capacity to utilize host protease TMPRSS2 for entry. Together, these data indicates that post-translational modification of Y674 impacts SARS-CoV-2 infection of the airway by disrupting both spike processing and host protease function. These results add to data demonstrating the importance of spike S1/S2 loop modifications to SARS-CoV-2 infection, pathogenesis, and transmission.

**Significance:** The S1/S2 loop of the SARS-CoV-2 spike protein distinguishes it from other sarbecoviruses and governs how efficiently the virus is activated for entry into host cells. While prior studies have disrupted this loop by deleting the furin cleavage site, shortening the loop, or mutating individual residues, each of these approaches focuses on the amino acid sequence itself. This study expands analysis to consider how post-translational modifications of amino acids within the loop contribute to infection. Mass spectrometry of purified virions identifies possible phosphorylation at Y674, a residue conserved across the Sarbecovirus family and maintained in all major SARS-CoV-2 variants. Our study finds that disrupting Y674 reduces spike processing on virions and diminishes usage of the host protease TMPRSS2, attenuating infection in human respiratory cells and in the airways of hamsters while not altering disease in the lung. Overall, the manuscript demonstrates that post-translational modification of the spike S1/S2 loop shapes SARS-CoV-2 entry and tropism. In addition, the work identifies a highly conserved residue playing a role in Sarbecovirus spike processing with implications as a target for coronavirus surveillance and therapeutic development.

## Introduction

Since the turn of the century, three coronaviruses (CoVs) have emerged and caused significant disease in the human population: Severe Acute Respiratory Syndrome Coronavirus (SARS-CoV, 2002), Middle East Respiratory Syndrome Coronavirus (MERS-CoV, 2012), and Severe Acute Respiratory Syndrome Coronavirus 2 (SARS-CoV-2, 2019)(1). SARS-CoV-2, the causative agent of CoVID-19, resulted in a large-scale pandemic that disrupted global public health and the world economy (2, 3). While effective vaccines, therapeutics, and infection induced immunity eventually resolved the pandemic, the repeated nature of CoV emergence highlights the need for continued study to predict, prevent, and mitigate future emergence events. With the broad and diverse range of CoV harbored in animals (4–8), efforts are needed not only to surveil but also understand key mechanisms of CoV infection to design better interventions and therapeutics.

While the pandemic emergency has been resolved, SARS-CoV-2 continues to circulate and evolve in the human population (9). New COVID-19 vaccines and therapeutics have blunted the worst elements of COVID-19 disease, but SARS-CoV-2 infections have produced novel variants that harbor mechanisms of enhanced immune evasion and viral fitness(10–13). Most of the changes in these SARS-CoV-2 variants have been linked with the spike protein receptor binding domain and increased capacity to evade host immunity (14, 15). Other changes since the original emergent SARS-CoV-2 strain have been associated with improved fitness and transmission in the variants (16–19). The later mutations, generally occurring in the terminal portion of the S1 domain of spike, largely impact spike processing and host protease utilization (14, 15). These results demonstrate that CoV spike processing, in addition to receptor binding, plays a critical role in SARS-CoV-2 infection and pathogenesis. Thus, to prepare and mitigate future CoV emergence events, we must build understanding of the elements that govern spike activation and fusion.

Unlike other sarbecoviruses, SARS-CoV-2 spike has two features in its S1/S2 loop that promote spike processing: an unusual furin cleavage site (FCS) and an extended S1/S2 loop (20, 21). Our group and others have previously shown that disruption of the FCS by either deletion or disruption results in attenuated SARS-CoV-2 infection, pathogenesis, and transmission (21–23). We subsequently showed that reducing the length of the S1/S2 loop resulted in similar attenuation, highlighting that both the FCS and extended loop were necessary for SARS-CoV-2 pathogenesis (20). As the pandemic continued, SARS-CoV-2 variants accumulated mutations impacting spike processing including mutations Y655H and P681H/R that improved spike processing compared to the original Wuhan strains(24). Our group recently showed that S1/S2 loop mutation N679K, conserved in every SARS-CoV-2 Omicron variant, reduced spike stability/expression and impacted both pathogenesis and transmission (12). Together, these results highlight the importance of spike processing on key aspects of SARS-CoV-2 infection, pathogenesis, and transmission.

Here, we seek further understanding of how CoV spike processing is impacted by mutations and modifications on the S1/S2 loop of spike. While deletion and disruption of key motifs like the FCS or the extended loop produced significant attenuation, single amino acid changes like N679K and P681H/R also had a major impact on SARS-CoV-2 infection and pathogenesis (12). With this in mind, we revisited analysis of post-translational modification of residues within the S1/S2 loop of the SARS-CoV-2 spike (20). Using mass spectrometry of purified virions, we had found that T678 in the spike S1/S2 loop is often glycosylated in the original emergent SARS-CoV-2 WA.1 strain . We subsequently generated a series of spike S1/S2 loop mutants that disrupted potential glycosylation sites (T676/T678) and found these mutants to be attenuated in host protease usage. These results highlighted that disruption of post-translational modifications in the S1/S2 loop could also shape SARS-CoV-2 infection.

In this study, we examine post-translation modifications and conservation of key residues in the S1/S2 loop. Using mass spectrometry data of both WT and mutant SARS-CoV-2, we identified post translational modifications (PTMs) on key residues within the S1/S2 loop. We subsequently focus study on phosphorylation at Y674 and generate a SARS-CoV-2 mutant (Y674F) which prevents this modification but maintains the amino acid structure. In vitro analysis of the Y674F mutant revealed attenuation in Calu3 cells. While in vivo disease was similar, we noted attenuation in viral replication in the upper airways as well as the large airways in the lung. Finally, we observed that loss of Y674 ablated spike processing on purified virions and its attenuation was likely due to less efficient usage of TMPRSS2. Overall, these results highlight the importance of Y674 in SARS-CoV-2 infection and its conservation may indicate a potential target for CoV surveillance and therapeutic targeting.

## Results

### Examine post-translational modification of residues in the SARS-CoV-2 S1/S2 loop

In this study, we focused on post-translational modifications (PTMs) to residues within the spike S1/S2 loop from purified WT and mutant SARS-CoV-2 virions (**Fig. 1A**). Using sucrose cushion purified virions, we previously performed LC-MS/MS and captured peptides across the spike protein from WT SARS-CoV-2, shortened loop mutant ΔQTQTN, and glycosylation mutants QTQVN and QVQVN . Here, we more closely examine other PTMs found in the mass spectrometry data occurring between spike residues 671 – 685, the amino acids comprising the S1/S2 loop prior to the canonical cleavage site at R685. Our prior study identified glycosylation of T678 in WT SARS-CoV-2 that was disrupted due to its absence (T678V) in the S1/S2 loop mutant viruses (20). We subsequently examined the adjacent spike peptide in the S1/S2 loop and identified four novel PTMs within residues 670-682 in either WT or SARS-CoV-2 S1/S2 loop mutants (**Fig. 1B, STable 1**). We noted glycosylation of S673 was observed in spike peptides in each of our SARS-CoV-2 purified virion preps and is consistent with findings from previously reported overexpression studies (25–27). We also observed unique PTMs in the S1/S2 loop mutants. For example, we observed glycosylation at T676 in a subset of spike peptides from the QTQVN mutant, a somewhat expected compensation due to the T678V substitution (**Fig. 1B**) (28, 29). Finally, we noted a novel phosphorylation at Y674 in the double glycosylation mutant (QVQVN). Together, these changes in the S1/S2 loop mutants suggested an effort to recapitulate the lost T678 glycosylation with alternative upstream PTMs.

**Figure 1.**
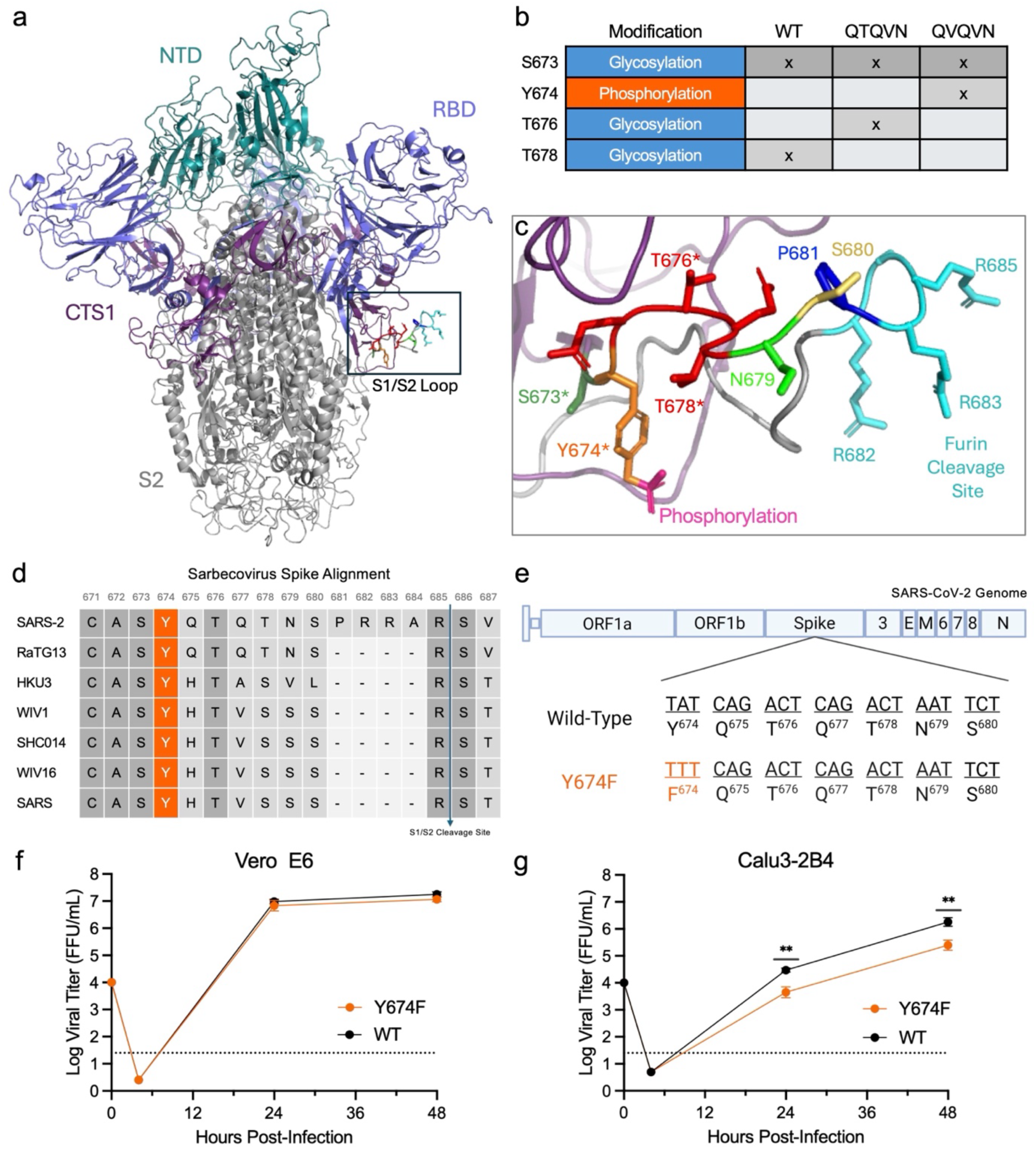
Mutation of spike residue Y674 attenuates SARS-CoV-2 replication. a) SARS-CoV-2 spike trimer structure showing the N-terminal domain (NTD, teal), receptor-binding domain (RBD-slate), C-terminal of S1 (CTS1-violet), and the S2 domain (grey). A S1/S2 loop from one of the spike trimers highlighted (black box). b) Table summarizing mass spectrometry findings of post-translational modification observed on spike residues between 670 – 685 in WT SARS-CoV-2 infection or S1/S2 loop mutants (QTQVN, QVQVN). c) Zoomed structure of the S1/S2 loop from a) showing phosphorylation of Y674 (residue-orange, phosphorylation-pink). Residues with * have observed post-translational modifications in one or more SARS-CoV-2 WT or mutants. d) Alignment of residues in the S1/S2 loop across representative members of the Sarbecovirus family. Numbers associated with position in SARS-CoV-2 WA.1 and highlights Y674 conservation across the family (orange). Other conserved residues represented by dark grey boxes. e) Schematic of SARS-CoV-2 spike with Y674F substitution identified. f) Viral titer from Vero E6 infected with WT (black) or Y674F (orange) SARS-CoV-2 at an MOI of 0.01 (n = 3 or 2). g) Viral titer from Calu-3 2B4 infected with WT or Y674F mutant SARS-CoV-2 at an MOI of 0.01 (n = 3). Data are mean ± SD. Statistical analysis was conducted using a two-tailed Student’s t test. **P ≤ 0.01. Experimental schematic in e) was created in BioRender. Menacherylab, T. (2026) https://BioRender.com/2n8n15r.

### Evaluate conserved and structural elements of spike S1/S2 loop residue Y674

We subsequently examined residue Y674 phosphorylation in the context of the spike structure and the S1/S2 loop. Ten amino acids from the canonical cleavage site, Y674 is a large, aromatic rings harboring amino acid located towards the start of the S1/S2 loop (**Fig. 1C**). Adjacent to established glycosylation sites at S683 and T678 (30, 31), steric hinderance of Y674 by these PTMs may prevent phosphorylation on WT SARS-CoV-2 spike (**Fig. 1C**). As such, Y674 phosphorylation on the QVQVN mutant may compensate for the loss of glycosylation at the adjacent T678 or T676. Alternatively, the two flanking glycosylations (T678, S673) may also obscure detection of the phosphorylation at Y674 on the WT spike. We note mutation of T678V and T676V in the QVQVN mutant disrupts the glycosylation sites and may have facilitated capturing the Y674 phosphorylation event (**Fig. 1B**). Based on its location in the S1/S2 loop, Y674 may play an important role in shaping SARS-CoV-2 infection and pathogenesis.

Having identified potential compensatory PTMs in the S1/S2 loop mutants, we next further evaluated spike residue Y674 in the light of its unique phosphorylation. We first evaluated conservation of Y674 across the Sarbecovirus family (**Fig. 1D**). We noted that Y674 is the last amino acid in a highly conserved region of spike between residues 659-674. Examining Sarbecovirus progenitor strain HKU3, epidemic SARS-CoV and SARS-CoV-2, and bat CoVs closely related to both epidemic strains (SHC014, WIV1, WIV16, RATG13), we found that amino acids 659-674 were 100% conserved across these sarbecoviruses (**Fig. 1D**). In contrast, the adjacent region of the S1/S2 loop has several amino acid differences across the viral family. In addition, using the UCSC Genome Browser (32), we observed that residue Y674 is completely conserved in all major SARS-CoV-2 variants ranging from the original Wuhan strain through Omicron variant XFG (https://genome.ucsc.edu/s/l/TSUbRIRoRiaZgP24GCG3dfdB). Overall, the results highlight Y674 as a stable and key residue in the S1/S2 loop across sarbecoviruses.

### Disruption of Y674 phosphorylation in SARS-CoV-2

To determine its role in SARS-CoV-2 infection, we used our reverse genetics system to disrupt Y674 phosphorylation in the SARS-CoV-2 WA.1 strain. We first considered an aggressive substitution to disrupt the amino acid function (Y674A). However, given its broad conservation across sarbecoviruses, we worried loss of the large aromatic residue may alter the shape of the S1/S2 loop (**Fig. 1C**). Instead, we focused on maintaining the loop structure but disrupting its phosphorylation capacity with substitution of phenylalanine (Y674F) (**Fig. 1E**). The Y674F mutation ablates phosphorylation by removing the free hydroxyl group sidechain while maintaining the aromatic ring structure, thus preserving the S1/S2 loop architecture. Using our established SARS-CoV-2 reverse genetic system (33, 34), we successfully recovered the Y674F mutant in the SARS-CoV-2 WA-1 backbone. The SARS-CoV-2 Y674F mutant had robust stock titers and showed no specific deficits in stock virus amplification or plaque morphology.

### Y674F mutant has attenuated viral replication in human airway cells

We next evaluated viral replication capacity of the Y674F mutant in our standard cell line models of SARS-CoV-2. We started with Vero E6 cells, an African green monkey kidney cells which lack a type I interferon response and are often used to amplify coronaviruses (35, 36). Following infection at MOI 0.01, we observed no deficit for the replication of Y674F mutant as compared to WT SARS-CoV-2 in Vero cells (**Fig. 1F**). These results are consistent with our findings with spike glycosylation mutants (QVQVN, QTQVN)(37) and suggest that the underlying mechanics of SARS-CoV-2 replication remain intact in the Y674F mutant. We subsequently evaluated Calu3 2B4 cells, a type I interferon competent human respiratory cell line commonly used in CoV research (38). In contrast to Vero E6, we observed that the Y674F mutant had a significant reduction in viral titer compared to WT SARS-CoV-2 (**Fig. 1G**). At both 24 and 48 hours post infection (hpi), the Y674F mutant had ∼1 log reduction in viral titer relative to WT control. Notably, these results differ from our findings with spike glycosylation mutants that had attenuated replication at 24 hpi but recovered by 48 hpi (37). Overall, the results indicate that disruption of Y674 phosphorylation capacity attenuated SARS-CoV-2 replication in human respiratory cells.

### Y674F attenuates viral replication in upper airway but not disease *in vivo*

Having established attenuation in human airway cells, we next sought to evaluate how the Y674F mutant impacts *in vivo* infection using the SARS-CoV-2 golden Syrian hamster model (39). Briefly, 3-4 weeks old male hamsters were intranasally infected with mock (PBS) or 10^5^ focus-forming units (FFUs) of either WT or Y674F mutant SARS-CoV-2 (**Fig. 2A**). Hamsters were subsequently observed for clinical disease over a 7-day time course with animals nasal washed and euthanized for tissue collection at days 2, 4, and 7 days post infection (dpi) (**Fig. 2A**). While mock infected animals gained weight, both WT (black) and Y674F mutant (orange) infection resulted in similar weight loss over the time course (**Fig. 2B**). For both infection groups, weight loss started at day 2 and peaked at ∼16-18% 6 dpi. Together, the results indicate that the Y674F mutant had no significant attenuation in clinical disease as compared to WT SARS-CoV-2.

**Figure 2.**
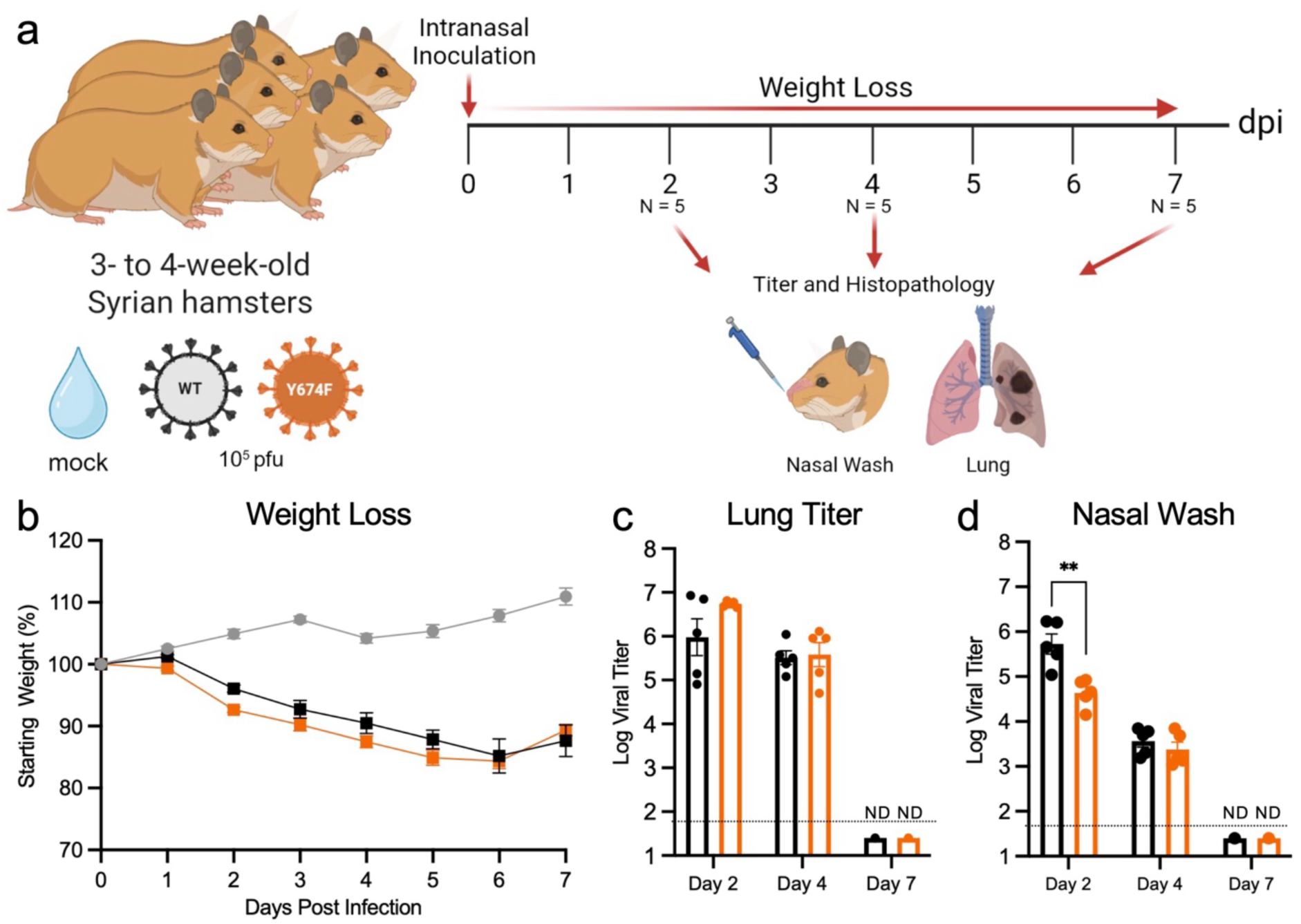
Y674F mutant attenuates viral replication in upper airway but not disease *in vivo*. a-b) Schematic of WT (black) or Y674F (orange) SARS-CoV-2 infection of golden Syrian hamsters. Three-to four-week-old male hamsters were infected intranasally with 10^5^ ffu of either virus and monitored for b) weight loss and disease for 7 days post infection. c-d) Viral titers were measured at day 2, 4, and 7 from c) infected lung or d) nasal wash. Data are representative of mean ± SEM. Statistical analysis was conducted using a two-tailed Student’s t test. **P ≤ 0.01. Experimental schematic in a) was created in BioRender. Menacherylab, T. (2026) https://BioRender.com/m1tlvll.

We next evaluated viral load in both the upper and lower respiratory tracts from hamsters following infection WT or Y674F mutant. We observed that the lungs of both WT and Y674F had robust titers at days 2 and 4 post infection (**Fig. 2C**). Viral loads of the mutant and WT were largely similar at both times with no significant differences observed. By day 7, viral clearance had occurred with no detectable virus observed in either group, consistent with our prior SARS-CoV-2 studies in hamsters (37, 40, 41). We subsequently evaluated nasal wash as a measure of upper airway replication (**Fig. 2D**). Examining day 2 nasal wash titers, we noted a significantly greater viral load in the WT infected hamster as compared to Y674F mutant. However, by day 4, no significant difference was observed between WT and Y647F mutant in the nasal titers. Similar to the lung, viral load was not detectable in nasal wash at day 7 post infection. Together, the results indicated that the Y674F mutant had a modest, but significant attenuation in the nasal wash at early times post infection. In contrast, the hamster lung was largely not impacted by the Y674F mutation in terms of viral load.

### Loss of Y674 has minimal impact on immune infiltrate and inflammation in the lung

To further evaluate disease and damage, mock, WT and Y674F mutant infected hamsters were evaluated for changes in lung histopathology. Lung sections were H&E stained and evaluated by a board-certified pathologist in a blinded manner as previously described (42). Consistent with previous studies (37, 40, 41), lung sections from day 2 in both WT and Y674F were largely unremarkable with minimal signs of disease or infiltration, similar to mock infected animals (**Fig. 3A-B**). However, by day 4, both WT and Y674F mutant infection had signs of severe pathology characterized by significant immune cell infiltration and inflammation (**Fig. 3A,C-D**). This lung damage advanced to severe lesions in both WT and Y674F mutants involving >70% of the lung tissue (**Fig. 3A, E-F**). Overall, the disease and damage at day 7 correspond to the more severe weight loss (**Fig. 2B**) and highlighted that the Y674F mutation had no major deficit in causing SARS-CoV-2 disease in the lung.

**Figure 3.**
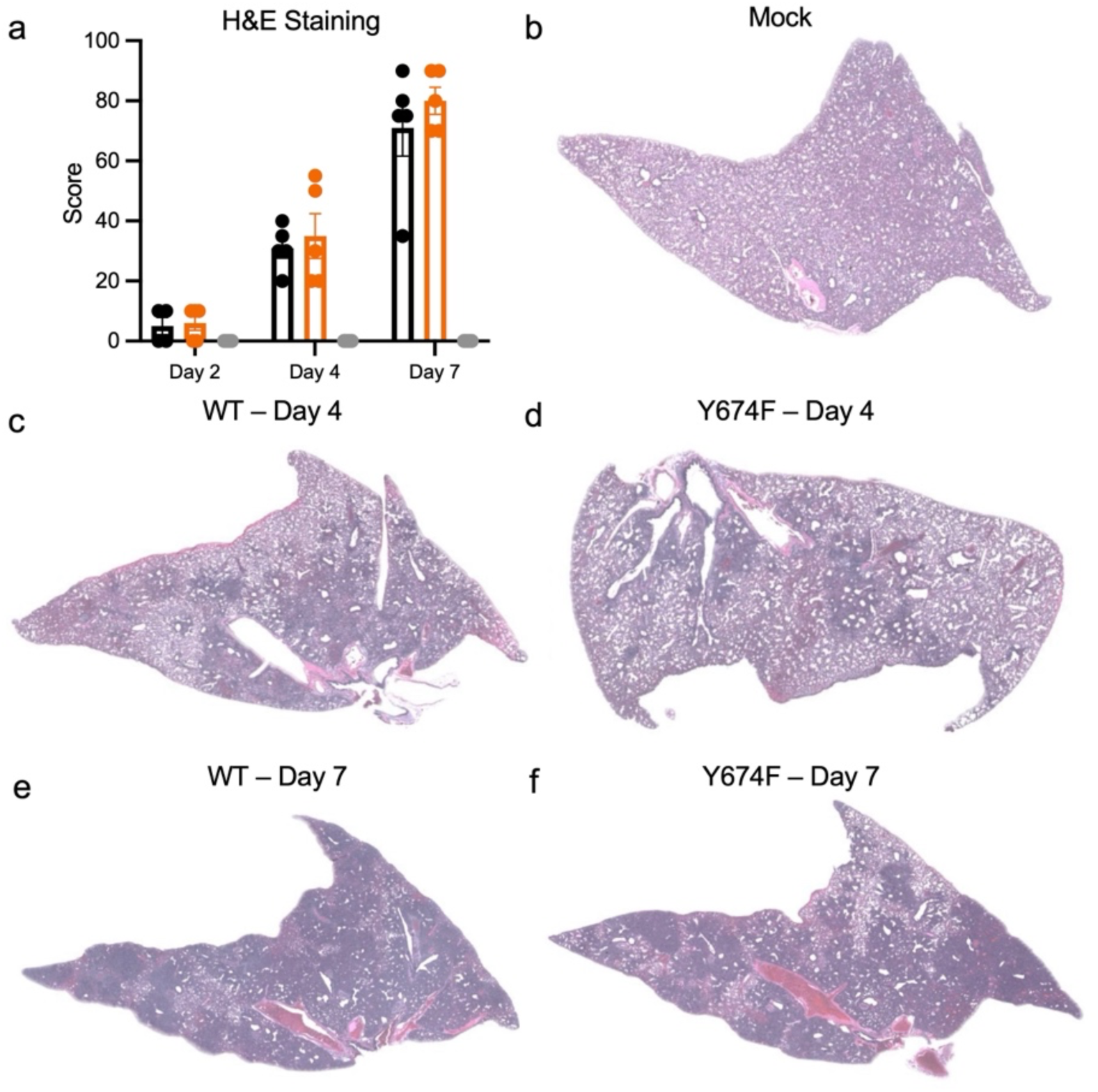
Loss of Y674 has minimal impact on lung inflammation. a) Lung histology sections from hamsters infected with WT (black), Y674F mutant (orange), or mock (grey) were H& E stained and scored for histopathological analysis at days 2, 4, and 7 post infection. Sections from individual animals were averaged and representing a single point within each group. Statistical analysis was conducted using a two-tailed Student’s t test and found no significant differences between the infected groups. b-f) Representative H&E staining of entire lung section of infected or mock infected hamsters from at designated times.

### Large airway antigen staining reduced in the Y674F mutant

Having observed no changes in immune infiltration or inflammation in the lung, we next evaluated viral distribution in lung. Prior studies with another SARS-CoV-2 S1/S2 loop mutant N679K had demonstrated a shift in antigen distribution with enhanced staining of large airways in the infected lungs (12). Here, we examined antigen distribution in the hamster lungs at days 2 and 4 post WT or Y674F infection. While no staining was observed in mock infected animals (**Fig. 4A & B**), we noted robust antigen distribution in WT infected animals at both day 2 and day 4 (**Fig. 4C & D**). SARS-CoV-2 antigen was observed in both airway and parenchyma of WT infected hamsters indicating broad distribution in the lung, consistent with prior studies (12, 37, 40–42). Similarly, Y674F also produced robust antigen staining at both days 2 and 4 (**Fig. 4E & F**). Scoring indicated that antigen staining of the Y674F mutant was largely similar to WT in both the parenchyma and overall staining (**Fig. 4G & H**). However, we noted attenuation of Y674F mutant staining in the airways at day 2 post infection (**Fig. 4I**). While equalized by day 4, the attenuation in antigen stain corresponded with reduced viral loads in the nasal washes of the Y674F mutant (**Fig. 2E**). Together, the results suggest that loss of Y674 phosphorylation reduces infection of airway cells following SARS-CoV-2 infection *in vivo*.

**Figure 4.**
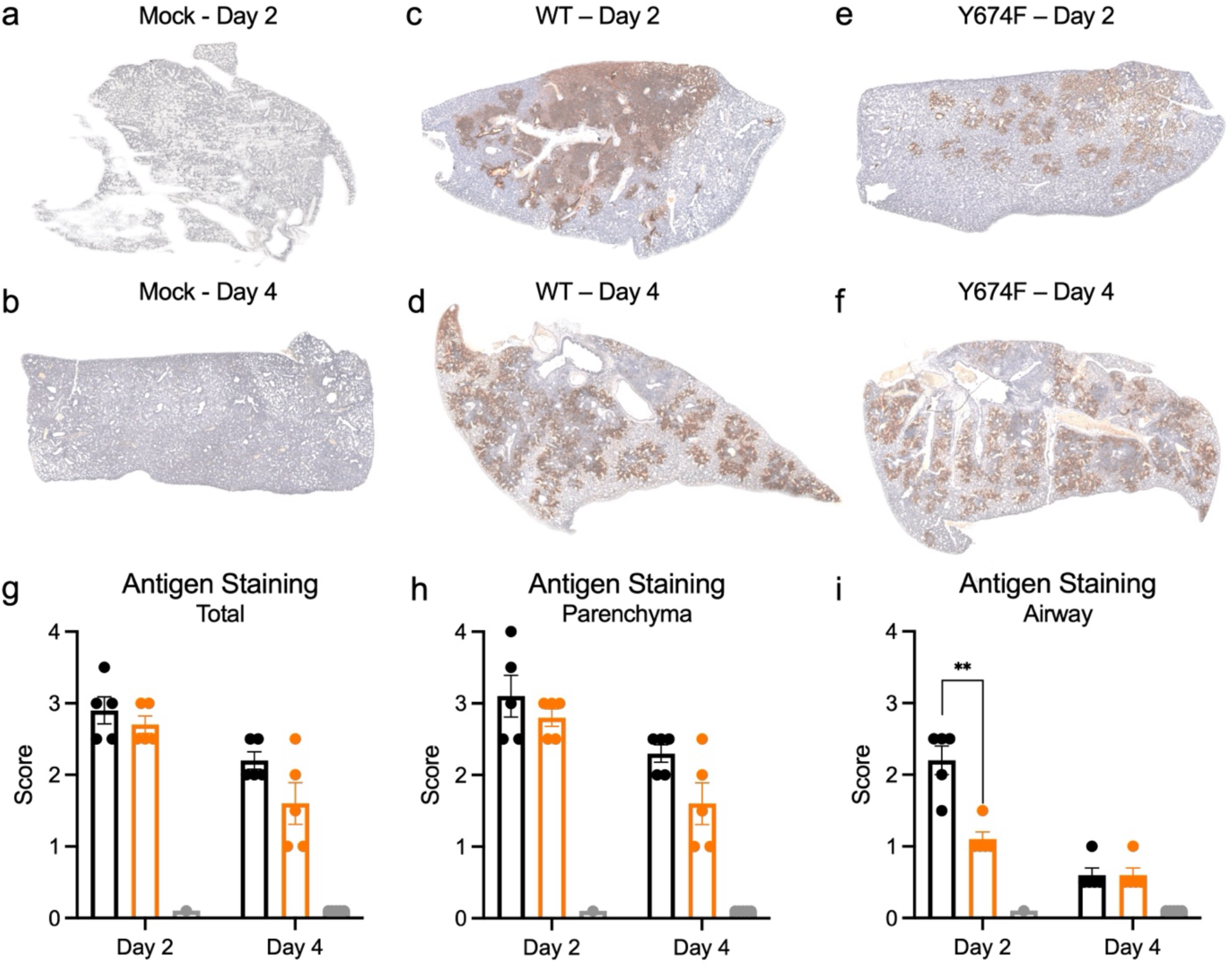
The Y674F has reduced antigen staining in the lungs . a-f) Nucleocapsid antigen staining of the lung sections from hamsters infected with a-b) mock c-d) WT or e-f) Y674F at day 2 or day 4 post infection. g-i) Antigen staining was scored for WT (black), Y674F (orange), or mock (grey) infected hamsters in a blinded manner for g) total distribution or location in the h) parenchyma or i) airway. Statistical analysis was conducted using a two-tailed Student’s t test. **P ≤ 0.01.

### Disruption of Y674 alters spike processing on virions

Spike-mediated entry requires sequential cleavage at the S1/S2 junction and the S2′ site to activate fusion and entry. The furin cleavage site in the S1/S2 loop of SARS-CoV-2 is known to facilitate spike processing on virions from full length into S1 and S2 subunits(43). Viruses that mutate the FCS or shorten the S1/S2 loop disrupt spike processing (37, 41). However, we also found that S1/S2 loop glycosylation mutants had no deficit in spike processing (37). Here, we examined spike processing on Y674F mutant virions after amplification on Vero E6 cells. Briefly, WT and Y674F SARS-CoV-2 were collected from infected Vero E6 cells at 24 hours post-inoculation, purified using ultracentrifugation sucrose cushion, inactivated, and blotted for spike and nucleocapsid expression (**Fig. 5A**). Normalizing protein levels to nucleocapsid, we observe that the Y674F mutant has a striking decrease in S1/S2 cleavage product (**Fig. 5B**). Quantifying the bands, WT virions have 65% of its spike processed to the S1/S2 product versus uncleaved full-length (∼35%) (**Fig. 5C**). In contrast, the Y674F mutant virions spike remains largely unprocessed with nearly all in full-length form (∼97%). Unlike the previously examined S1/S2 loop mutants (37), loss of Y674 phosphorylation appears to limit spike processing on SARS-CoV-2 virions.

**Figure 5.**
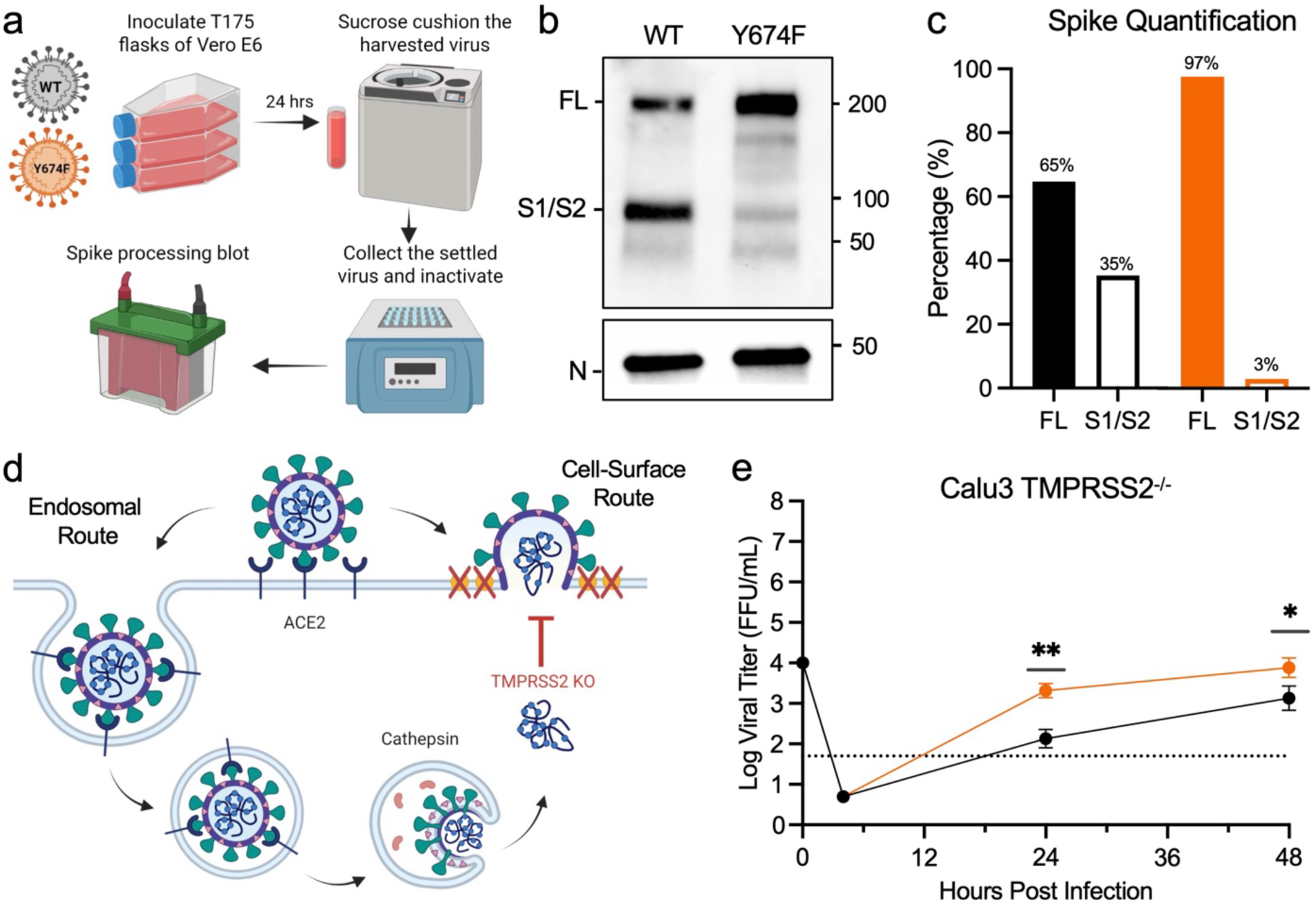
Loss of Y674 impairs spike processing and alters host protease usage. a) Schematic of SARS-CoV-2 virion collection vial sucrose cushion. b) Sucrose cushion purified WT and Y674F virions from infected Vero E6 cells were probed with α-Spike and α-Nucleocapsid (N) antibodies by Western blot. Full-length spike (FL), /S2 cleavage, and N are indicated. c) Quantification of densitometry for FL (closed bar) and S1/S2 (open bar) are shown for WT (black) and Y674F mutant (orange). d) Schematic of SARS-CoV-2 entry pathways distinguishing endosomal and cell surface routes. Diagram shows disruption of cell surface entry through knockout of TMPRSS2. e) Viral titer from Calu-3 TMPRSS2 knock-out cells infected with WT (black) or Y674F mutant (orange) SARS-CoV-2 at an MOI of 0.01 (n = 3). Statistical analysis was conducted using a two-tailed Student’s t test. *P ≤ 0.05; **P ≤ 0.01. Schematic in a) and d) were created in BioRender. Menacherylab, T. (2026) https://BioRender.com/nj48w4b; https://BioRender.com/2qvdseh.

### Loss of TMPRSS2 mediated entry does not impact Y674F mutant

SARS-CoV-2 utilizes two pathways for entry into host cells: 1) the endosomal route mediated by spike activation by host cathepsins or 2) the cell surface route mediated by host serine proteases like TMPRSS2 (**Fig. 5D**). Prior studies had determined that entry of SARS-CoV-2 into human Calu3 cells is driven primarily by TMPRRS2 (44, 45). In addition, we had shown that attenuation of a furin cleavage site mutant was ablated in Calu3 cells lacking TMPRSS2 (41). Since Y674F had significant attenuation in Calu3 cells (**Fig. 1G**), we subsequently evaluated the mutant in Calu3 cell lacking TMPRSS2. Consistent with prior studies, loss of TMPRSS2 resulted in reduced viral replication of WT SARS-CoV-2 (**Fig. 5E**). WT SARS-CoV-2 replication in TMPRSS2^-/-^ Calu3 cells had reductions of 2.3 and 3.1 log titer at 24 and 48 hpi compared to control Calu3 (**Fig. 1G**). In contrast, the Y674F mutant had similar titers with the Calu3 TMPRSS2^-/-^ cells having a modest 0.3 log reduction in viral titer compared to control Calu3 24 hpi (**Fig. 1G**). This TMPRSS2 based attenuation of Y674F expands to 1.5 log reduction at 48 hpi. However, these results demonstrate reduced capacity of the Y674F mutant to utilize TMPRSS2 compared to WT

## Discussion

While host receptor binding plays a critical role in CoV infection, the impact of spike activation and entry cannot be discounted. Our prior work highlighted the critical importance of the FCS and extended S1/S2 loop in SARS-CoV-2 infection, pathogenesis, and transmission (12, 20, 22, 23). Here, we demonstrate that post-translational modifications on the S1/S2 loop can also play an important role in the dynamics of SARS-CoV-2 infection. Using reverse genetics, we disrupted a potential phosphorylation site at Y674 while maintaining the amino acid structure in S1/S2 loop of the SARS-CoV-2 spike (**Fig. 1E**). The resulting spike Y674F mutant was attenuated in human respiratory cells *in vitro* and in the airways *in vivo* (**Fig. 1G**, **Fig. 2D**, **Fig. 4I**). Loss of the capacity to phosphorylate the conserved Y674 resulted in reduced spike processing on purified virions as well as reduced ability to utilize TMPRSS2 for infection (**Fig. 5**). Overall, the data highlights that post-translational modification of the S1/S2 loop of spike plays a critical role in shaping SARS-CoV-2 infection tropism and has implications for pathogenesis and transmission.

By examining virion PTMs using mass spectrometry, we have identified another landscape that shapes CoV spike processing and host protease usage. Our initial examination of purified virions from WT SARS-CoV-2 WA.1 revealed glycosylation in the S1/S2 loop at both T673 and T678 (**Fig. 1B**) (37). However, only after examining the glycosylation mutant that disrupted both T678 and T676 were we able to observe phosphorylation of Y674. This left the possibility that Y674 gets phosphorylated in compensation for the loss of flanking glycosylations at T678 or T676. Alternatively, glycosylation at T673 and T678 may have obscured Y674 phosphorylation in WT spike. While alternative mass spectrometry approaches could resolve this question, we instead used reverse genetics to ablate phosphorylation with a Y674F substitution. Absent the hydroxyl group side chain, phenylalanine at position 674 maintains the amino acid structure but disrupts phosphorylation capacity. The resulting Y674F mutant had robust replication in Vero cells, but attenuation in Calu3 cells and in the airway *in vivo* (**Fig. 1G, 2D, 4I**). Loss of Y674 disrupted spike cleavage on purified SARS-CoV-2 virions and diminished the capacity to use TMPRSS2 for entry (**Fig. 5**). Importantly, Y674 is conserved across the sarbecoviruses suggesting that that phosphorylation of this residue may play a critical role in spike processing and host protease usage across the entire family (**Fig. 1D**). While requiring further examination, these results may provide a new, highly conserved residue within Sarbecovirus spike to target for therapeutic treatment.

Our findings further demonstrate the importance of PTMs on the S1/S2 loop of spike for CoV infection, pathogenesis, and transmission. Here, the loss of Y674 phosphorylation capacity ablates spike processing on virions and disrupts the use of TMPRSS2 (**Fig. 5**). Prior S1/S2 glycosylation mutants (QVQVN, QTQVN) also had disrupted TMPRSS2 utilization, although infection was restored at late times unlike in the Y674F mutant (37). Our studies with a SARS-CoV-2 N679K mutant disrupted the amount of spike on the virion surface which may be due to a longer polybasic site; alternatively, N679K may serve as a potential ubiquitination site that causes the loss of spike on virions (12). Notably, N679K also changed the dynamics of transmission in the hamster model, shifting replication advantage to the upper airway. While not tested for either Y674F or the glycosylation mutants, PTM changes in the loop may also impact transmission. For each of the mutations in the S1/S2 loop described above, the alteration of the PTM changed spike processing, spike stability, or host protease usage, all key elements in SARS-CoV-2 entry and infection. Importantly, other conserved PTM sites like S673 have not yet been evaluated for their impact on SARS-CoV-2 infection and pathogenesis. In addition, with the continued evolution in SARS-CoV-2 variants of concern, numerous groups have identified broad changes in spike PTMs between the original SARS-CoV-2 Wuhan strains and Omicron lineage variants (46–48). These changes open the possibility that mutations that disrupt or produce PTMs could reshape critical aspects of SARS-CoV-2 infection. Beyond SARS-CoV-2, other epidemic CoVs like MERS-CoV or zoonotic strains may also have critical PTMs that impact infection and that may vary over time. By understanding how these PTMs impact infection and entry, we have the opportunity to develop surveillance approaches as well as target these PTMs with therapeutic approaches.

Overall, this study demonstrates disruption of phosphorylation capacity at Y674 in the S1/S2 loop of SARS-CoV-2 spike reduces spike processing on virions, disrupts host protease usage, and results in attenuated infection in airways. With its broad conservation across the Sarbecovirus family, our findings potentially identify a key residue to target with therapeutic approaches. In addition, the work combines with our prior studies to highlight the importance of PTMs in SARS-CoV-2 spike activation and host protease usage. Understanding these mechanisms has potential implications for not only sarbecoviruses, but can inform efforts to identify, prevent, and mitigate future CoV emergence events.

## MATERIALS AND METHODS

### Cells

Vero E6 cells were grown in Dulbecco modified Eagle medium (DMEM; Gibco #11965-092) supplemented with 10% fetal bovine serum 9FBS) (HyClone #SH30071.03) and 1% antibiotic-antimycotic (Gibco #5240062). Calu-3 2B4 cells were grown in DMEM supplemented with 10% defined FBS (HyClone #SH30070.03), 1% antibiotic-antimycotic, and 1 mg/mL sodium pyruvate. Human TMPRSS2 knockout Calu-3 cells (Abcam #273734) were grown in DMEM supplemented with 20% defined FBS (HyClone #SH30070.03), 1% antibiotic-antimycotic, 1% non-essential amino acid solution (NEAA; Gibco #11140050), and 1 mg/mL sodium pyruvate.

### Viruses

The recombinant WT and mutant SARS-CoV-2 virus sequences are based on the USA-WA1/2020 isolate sequence provided by the World Reference Center for Emerging Viruses and Arboviruses, which was originally obtained from the US Centers for Disease Control and Prevention (49). Wild-type and mutant SARS-CoV-2 were generated using standard cloning techniques and the reverse genetics system as previously described (33, 34) and propagated on Vero E6 cells. Mutations in the Y674F mutants were verified with Sanger sequencing. Recovered mutant virus was further sequenced with Tiled Clickseq NGS to confirm the maintenance of nucleotide mutations up to P2 (50). Infectious titers were measured by focus-forming assay. All experiments involving infectious virus were conducted at the University of Texas Medical Branch (UTMB) in an approved biosafety level 3 laboratory with routine medical monitoring of staff.

### Structure modeling

Structural models were generated using SWISS-Model to generate homology models for WT SARS-CoV-2 spike protein on the basis of the SARS-CoV-1 trimer structure (Protein Data Bank code 6ACD) as previously described (22). Homology models were visualized and manipulated in PyMOL (version 3.1.8).

### Nanoflow-LC-MS/MS Glycopeptide Mapping

Sucrose cushion-purified SARS-CoV-2 virion samples for WT and S1/S2 loop mutants were prepared for LC-MS/MS analysis and processed as previously described (19). The mass spectrometry proteomics data are available in the ProteomeXchange Consortium via the PRIDE partner repository with the dataset identifier PXD034332 (33) and 10.6019/PXD034332 (34)

### *In vitro* infection

Viral infections in Vero E6, Calu-3 2B4 and Calu-3 TMPRS2 KO cells were performed as previously described in (33, 51). Briefly, cells were washed with phosphate buffered saline (PBS) and infected with WT or mutant SARS-CoV-2 at an MOI of 0.01 for 45 min at 37 °C. Following absorption, cells were washed three times with PBS and fresh growth media was added. Three or more biological replicates were collected at each time point.

### Focus-forming assay

Focus-forming assays were performed as previously described (17). Briefly, Vero E6 cells were seeded in 96-well plates to be 100% confluent. Samples were 10-fold serially diluted in serum-free media, and 20 µL was used to infect cells. Cells were incubated for 45 min at 37°C with 5% CO2 before 100 µL of 0.85% methylcellulose overlay was added. Cells were incubated for 24 h at 37°C with 5% CO2. After incubation, the overlay was removed, cells were washed three times with PBS before being fixed, and the virus was inactivated by 10% formalin for 30 min at room temperature. Cells were then permeabilized and blocked with 0.1% saponin/0.1% BSA in PBS before being incubated with α-SARS-CoV-2 Nucleocapsid primary antibody (Cell Signaling Technology) at 1:1,000 in permeabilization/blocking buffer overnight at 4°C. Cells are then washed three times with PBS before being incubated with Alexa FluorTM 555-conjugated α-mouse secondary antibody (Invitrogen #A28180) at 1:2,000 in permeabilization/blocking buffer for 1 h at room temperature. Cells were washed three times with PBS. Fluorescent foci images were captured using a Cytation 7 cell imaging multi-mode reader (BioTek), and foci were counted manually.

### Virion purification and Western blotting

Vero E6 cells were infected with WT or mutant SARS-CoV-2 at an MOI of 0.01. Supernatant was harvested 24 hpi and clarified by low-speed centrifugation. Virus particles were then pelleted by ultracentrifugation through a 20% sucrose cushion at 26,000 rpm for 3 h using a Beckman SW28 rotor. Pellets were resuspended in 2× Laemmli buffer to obtain protein lysates. Relative viral protein levels were determined by sodium dodecyl sulfate-polyacrylamide gel electrophoresis (SDS-PAGE) followed by Western blot analysis as previously described^1,2^. In brief, sucrose-purified WT and mutant SARS-CoV-2 virions were inactivated by boiling in Laemmli buffer. Samples were loaded in equal volumes into 4%–20% Mini-PROTEAN TGX Gels (Bio-Rad #4561093) and electrophoresed by SDS-PAGE. Protein was transferred to polyvinylidene difluoride membranes. Membranes were probed with SARS-CoV S-specific antibodies (Novus Biologicals #NB100-56578) and followed by horseradish peroxidase (HRP)-conjugated anti-rabbit antibody (Cell Signaling Technology #7074). Membranes were stripped and reprobed with SARS-CoV N-specific antibodies (Novus Biologicals #NB100-56576) and the HRP-conjugated anti-rabbit secondary IgG to measure loading. Signal was developed using Clarity Western ECL substrate (Bio-Rad #1705060) or Clarity Max Western ECL substrate (Bio-Rad #1705062) and imaged on a ChemiDoc MP System (Bio-Rad #12003154). Densitometry was performed using ImageLab 6.0.1 (Bio-Rad #2012931).

### Hamster infection studies

Male golden Syrian hamsters (3–4 weeks old) were purchased from Envigo (HsdHan:AURA strain). All studies were conducted under a protocol approved by the UTMB Institutional Animal Care and Use Committee and complied with USDA guidelines in a laboratory accredited by the Association for Assessment and Accreditation of Laboratory Animal Care. Procedures involving infectious SARS-CoV-2 were performed in the Galveston National Laboratory ABSL3 facility. The ABSL-3 room was kept between 68–74 degrees Fahrenheit with 30–60% and 12h-12h light cycles (6AM-6PM). Hamsters were intranasally inoculated with 10^5^ pfu of WT or Y674F SARS-CoV-2 in 100 μL. Infected hamsters were weighed and monitored for illness over 7 days. Hamsters were anesthetized with isoflurane (Henry Schein Animal Health), and nasal washes were collected with 400 μL of PBS on endpoint days (2, 4, and 7 dpi). Hamsters were euthanized by CO2 for organ collection. Nasal wash and lungs were collected to measure viral titer. The left lungs were collected for histopathology.

### Histology

Histopathology was performed as previously described (52, 53). Briefly, the left lungs were harvested from hamsters and fixed in 10% buffered formalin solution for at least 7 days. Fixed tissue was then embedded in paraffin, cut into 5 µm sections, and stained with hematoxylin and eosin on a SAKURA VIP6 processor by the University of Texas Medical Branch Surgical Pathology Laboratory. H&E staining was performed by the University of Texas Medical Branch Histology Laboratory and then analyzed and scored by a blinded pathologist.

### Immunohistochemistry

Immunostaining was conducted as previously described (54, 55). Briefly, fixed and paraffin-embedded left lung lobes from hamsters were cut into 5 µm sections and mounted onto slides by the University of Texas Medical Branch Surgical Pathology Laboratory. Paraffin-embedded sections were warmed at 56°C for 10 min, deparaffinized with xylene (3× 5-min washes) and graded ethanol (3× 100% 5-min washes and 1× 95% 5-min wash), and rehydrated in distilled water. After rehydration, antigen retrieval was performed by steaming slides in antigen retrieval solution (10 mM sodium citrate and 0.05% Tween-20, pH 6) for 40 min (antigen retrieval solution was boiled in a microwave, slides were added to the boiling solution, and incubated in a steamer). Slides were then cooled under running tap water for 10 min. Endogenous peroxidases were quenched by incubating slides in TBS with 0.3% H2O2 for 15 min followed by 2× 5 min washes in 0.05% TBST. Endogenous avidin and biotin were blocked by sequential 15-min incubations with avidin and biotin blocking solutions [Abcam #ab64212], each followed by 2× 5-min washes in TBST. Sections were blocked with 10% normal goat serum in BSA diluent (1% BSA in 0.05% TBST) for 30 min at room temperature. Sections were incubated with primary anti-N antibody (Sino #40143-R001) at 1:1,000 in BSA diluent overnight at 4°C. Following overnight primary antibody incubation, sections were washed 3× for 5 min in TBST. Sections were incubated in biotinylated anti-rabbit IgG secondary antibody (Vector Laboratories #BA-1000) at 1:200 in BSA diluent for 30 min at room temperature. Following secondary antibody incubation, sections were washed 3× for 5 min in TBST. Sections were then incubated in streptavidin-HRP (Vector Laboratories #SA-5004) at 1:200 in BSA diluent for 30 min at room temperature, followed by 3× 5-min washes in TBST. To visualize the antigen, sections were incubated in ImmPACT NovaRED (Vector Laboratories #SK-4805) for 3 min at room temperature before being rinsed with TBST to stop the reaction, followed by 1× 5-min wash in distilled water. Sections were incubated in hematoxylin for 5 min at room temperature to counterstain before rinsing in tap water for 2 min to stop the reaction. Sections were dehydrated by briefly passing them through the previous xylene and graded ethanol baths in reverse order before being mounted with coverslips using Permount (Fisher Scientific #SP15-500).

## Acknowledgments

Research was supported by grants from NIAID of the NIH to (R01-AI153602, R01-AI200317, and R21-AI145400 to V.D.M.). Research was also supported by STARs Award provided by the University of Texas System and Burroughs Welcome Fund Investigators in Pathogenesis grant to V.D.M. Data Acquisition award provided by the Institute for Human Infections and Immunity at UTMB to M.N.V. Trainee funding provided by NIAID of the NIH to M.N.V. (T32-AI060549), R.E.A. (T32-AI007526) and McLaughlin Fellowship to Y.A. (88208). Experiments and analysis were provided by the Mass Spectrometry Facility at the University of Texas Medical Branch (https://www.utmb.edu/MSF) and is funded in part by CPRIT RP250644 (WKR).

## Author contributions

Conceptualization: WMM, ALM, MNV, VDM

Formal analysis: WMM, ALM, MNV, KGL, YA, LKE, YZ, DHW, WKR VDM

Funding acquisition: MNV, YA, REA, WKR, VDM Investigation: WMM, ALM, MNV, KGL, YPA, LKE, DHW, VDM Methodology: WMM, ALM, MNV, YZ, KGL, VDM

Project Administration: WMM, MNV, VDM Supervision: MSS, BAJ, VDM

Visualization: WMM, ALM, MNV, DHW, VDM Writing – original draft: WMM, VDM

Writing – review and editing: WMM, VDM

## Supplemental Information

## Supplemental Information

**Supplemental Table S1.**
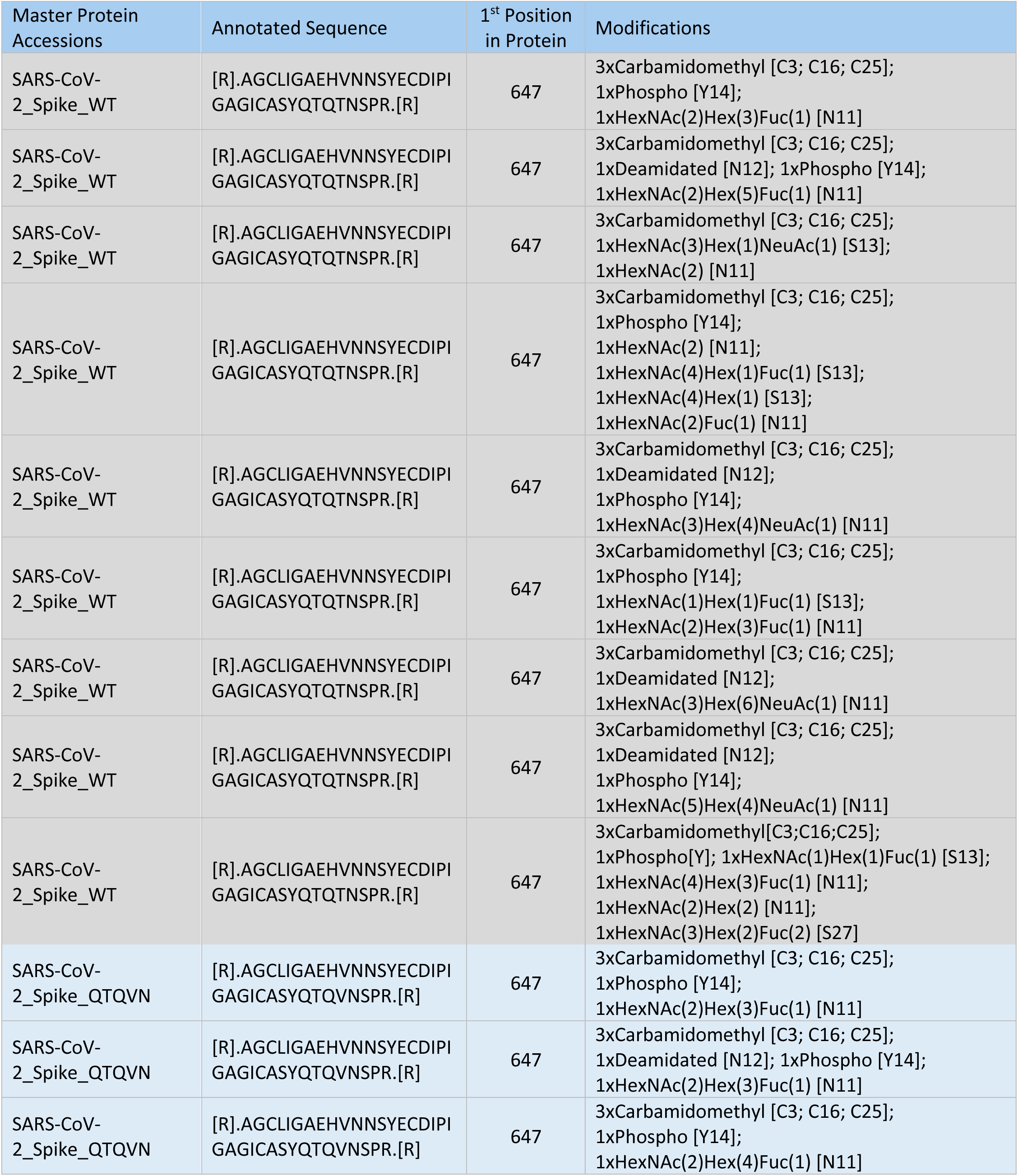
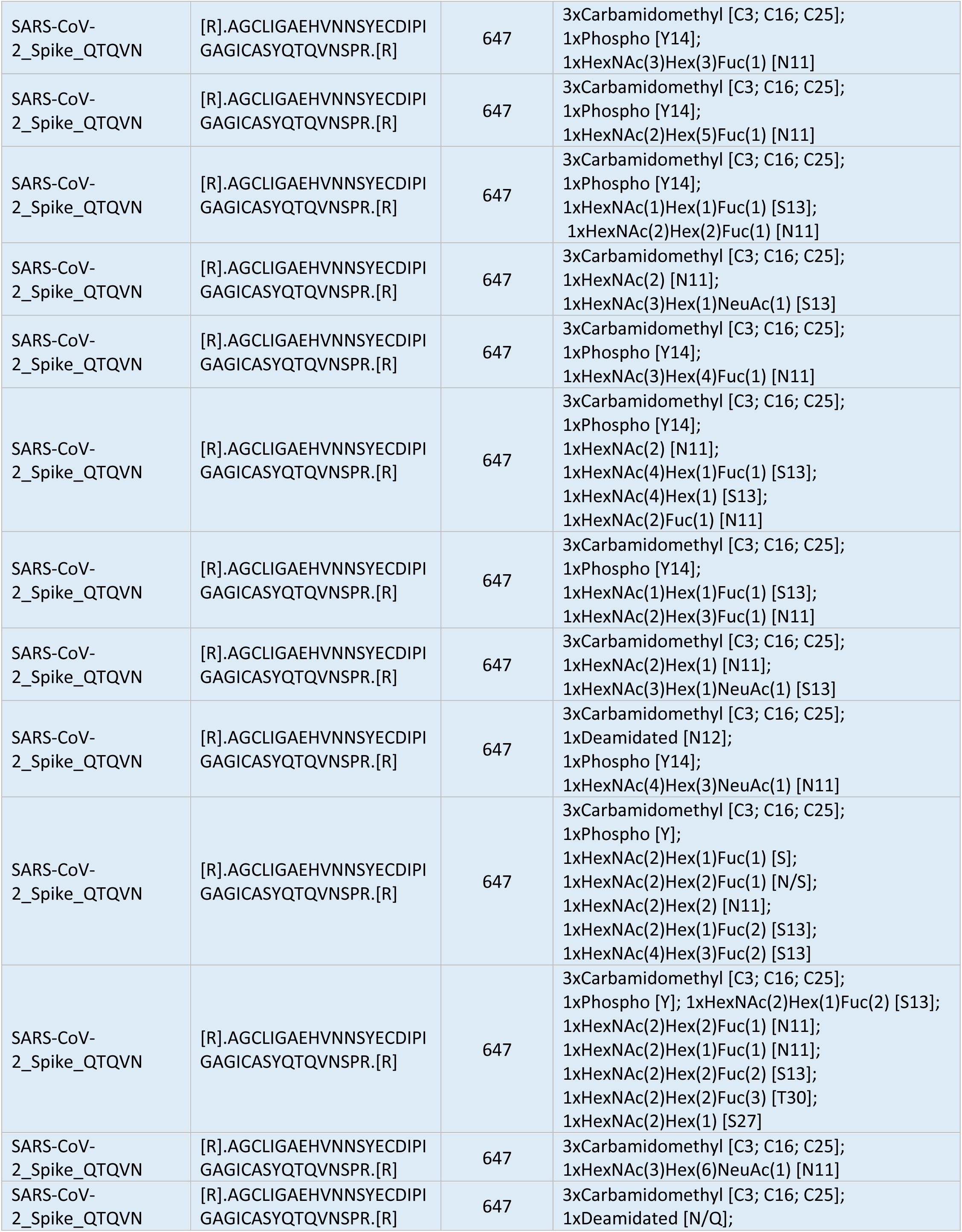
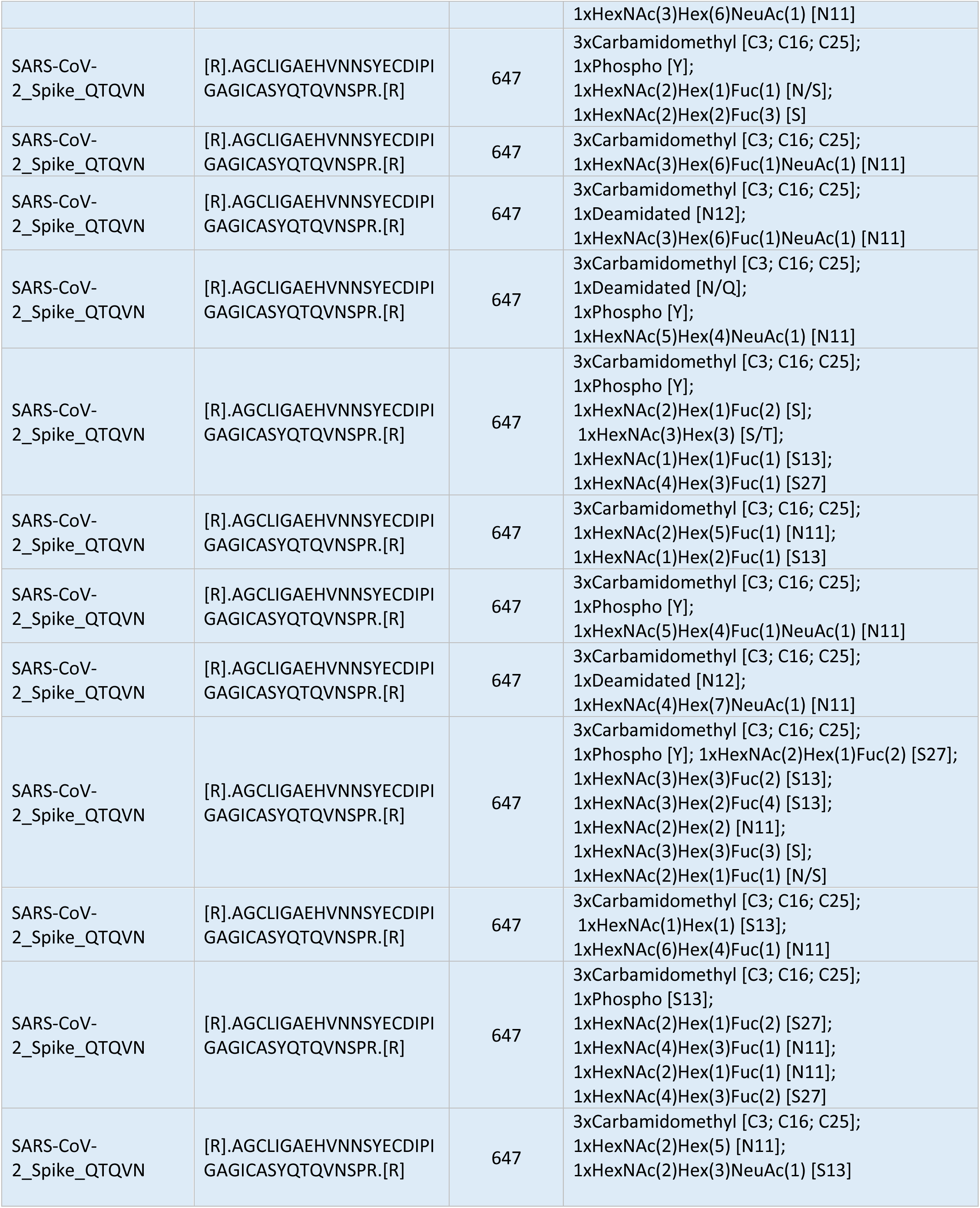
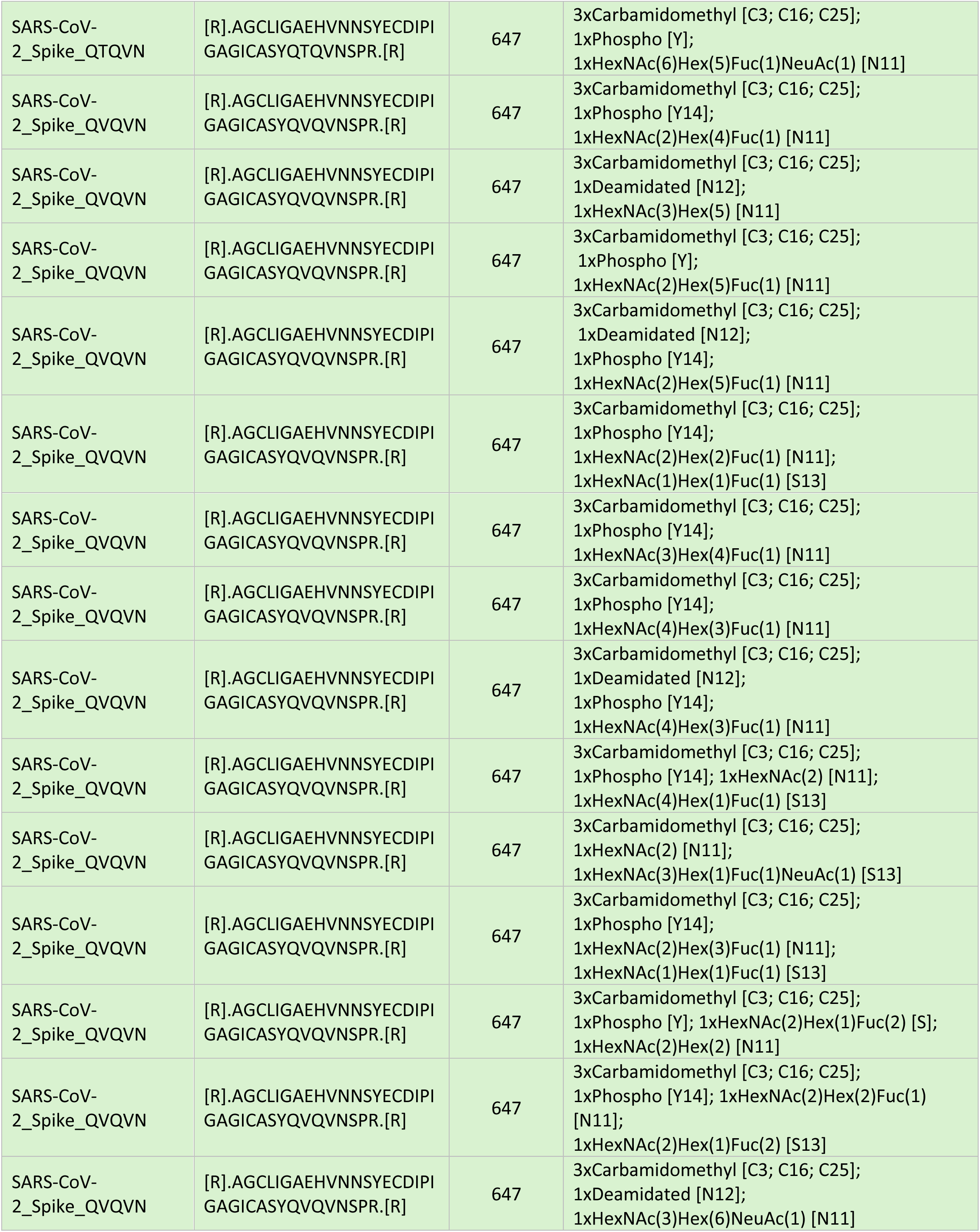
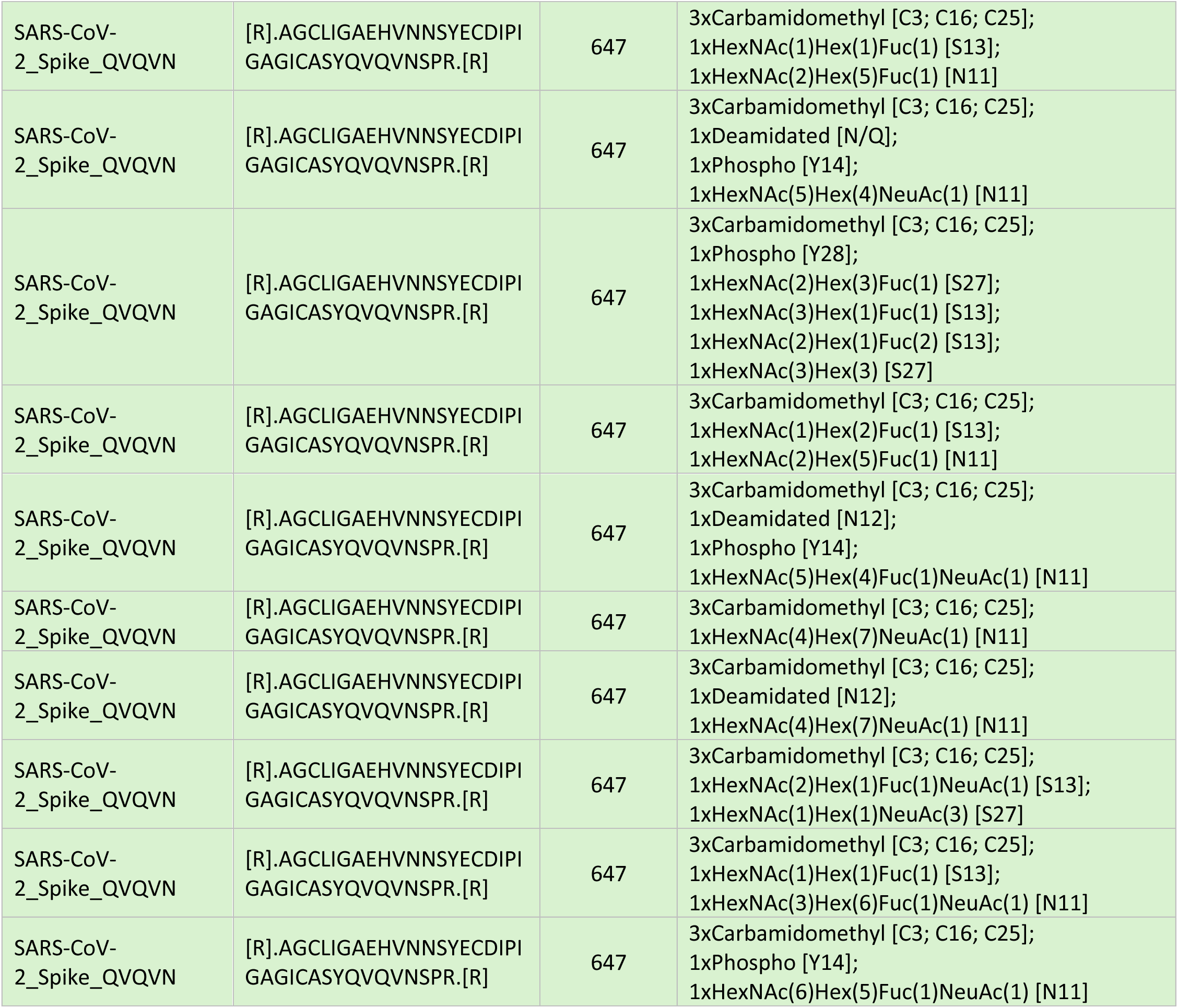
Quantification of Nanoflow-LC-MS/MS analysis of SARS-CoV-2 WT and glycosylation mutants. Modifications of peptide spanning Spike 647-682 for WT (grey), QTQVN (blue) and QVQVN (green).

